# Administration of the glutamate-modulating drug riluzole after stress reverses its delayed effects on the amygdala

**DOI:** 10.1101/2023.03.08.531819

**Authors:** Siddhartha Datta, Zubin Rashid, Saptarnab Naskar, Sumantra Chattarji

**Author notes:** **Corresponding Author:** *Prof. Sumantra Chattarji; E-Mail:. Equal contribution (Siddhartha Datta, Zubin Rashid). **Authors’ contributions:** S.D., Z.R., S.N., and S.C. designed the study. S.D., and Z.R. collected the data. S.D., Z.R., and S.N. compiled and analyzed the data. Z.R., S.N., and S.C. wrote the manuscript. S.C. supervised the study, provided funding and resources for the study, and edited the manuscript. **Competing interest statement:** The authors declare no competing financial interest.

## Abstract

Extracellular glutamate levels are elevated across brain regions immediately after stress. Despite sharing common features in their genesis, the patterns of stress-induced plasticity that eventually take shape are strikingly different between these brain areas. While stress impairs structure and function in the hippocampus, it has the opposite effect on the amygdala. Riluzole, an FDA-approved drug known to modulate glutamate release and facilitate glutamate clearance, prevents stress-induced deficits in the hippocampus. But, whether the same drug is also effective in countering the opposite effects of stress in the amygdala remains unexplored. We addressed this question by using a rat model wherein even a single 2-hour acute immobilization stress causes a delayed build-up, 10 days later, in anxiety-like behavior, alongside stronger excitatory synaptic connectivity in the basolateral amygdala (BLA). This temporal profile – several days separating the acute stressor and its delayed impact – allowed us to test if these effects can be reversed by administering riluzole in the drinking water *after* acute stress. Post-stress riluzole not only prevented the delayed increase in anxiety-like behavior on the elevated plus-maze, but also reversed the increase in spine-density on BLA neurons 10 days later. Further, stress-induced increase in the frequency of miniature excitatory postsynaptic currents recorded in BLA slices, 10 days later, was reversed by the same post-stress riluzole administration. Together, these findings underscore the importance of therapeutic strategies, aimed at glutamate uptake and modulation, in correcting the delayed behavioral, physiological, and morphological effects of stress on the amygdala.

**Significance statement:** Stress disorders are characterized by impaired cognitive function alongside enhanced emotionality. Consistent with this, the same stress elicits contrasting effects in the rodent hippocampus versus amygdala. This poses a therapeutic challenge – the same pharmacological intervention against stress has to counter these opposite effects. Yet, the immediate consequence of stress – enhanced extracellular glutamate – is similar across these two areas. To target this common feature, we treated rats with riluzole, a drug that prevents stress-induced glutamate surge. Although the drug was administered after the end of stress, it reversed its delayed impact on amygdalar structure and function. Since riluzole also enhances glutamate-uptake through glial-transporters and is approved for human use, these results highlight the importance of therapeutic strategies focused on neuron-astrocyte interactions.

## Introduction

The amygdala, a brain structure that is necessary for the encoding of emotionally salient memories, is susceptible to stressful experiences (1,2). Accumulating evidence from animal models has showed that stress leads to significant changes in the structure and function of amygdalar neurons, including growth of dendrites and spines, and increased transmission and plasticity at excitatory synapses (3,4). Stress also enhances conditioned fear and anxiety-like behavior (1,5-8). Together, these findings on stress-induced plasticity in the amygdala offer insights into the exaggerated affective symptoms seen in stress-related neuropsychiatric disorders (3,4,9-12). Increase in extracellular glutamate levels and glutamatergic transmission in the amygdala is one of the immediate consequences of stress (13,14). Hence, in this study we test the hypothesis that pharmacological attenuation of post-stress elevation in glutamate levels and glutamatergic transmission in the amygdala should reverse the delayed cellular and behavioral effects of stress. Further, instead of pharmacological interventions before or during stress, a clinically realistic paradigm would entail administering drugs *after* the stressful experience. It is in this context that a rodent model of acute immobilization stress offers an important advantage due to the unique temporal profile of changes it triggers in the amygdala (3,15-17). Specifically, a single 2-hour episode of immobilization stress leads to delayed morphological and physiological strengthening of synaptic connectivity in the basolateral amygdala (BLA), along with enhanced anxiety-like behavior, 10 days later (15-17). Thus, this rat model presents a window for post-stress intervention to prevent the immediate stress-induced increase of glutamate and then test if the gradual build-up in neuronal and behavioral changes can be blocked 10 days after acute stress.

Riluzole, a compound that enhances glutamate uptake, modulates glutamate release, and inhibits post-synaptic glutamate receptors (18-23), offers several advantages as a post-stress pharmacological intervention. First, this drug is approved for use in patients of amyotrophic lateral sclerosis (ALS) (24-27) because it counteracts glutamate excitotoxicity (28) by enhancing glutamate uptake mediated by the astrocytic transporters GLT-1 and GLAST (19,20), as well as attenuating presynaptic glutamate release (22). Further, riluzole is soluble in drinking water and hence bypasses the need for stressful interventions such as intraperitoneal injections or oral gavage in rodents (29,30). Interestingly, riluzole administered before or during chronic stress has been shown to block its effects in the hippocampus and prefrontal cortex (PFC) (31-34). However, nothing is known about the efficacy of post-stress riluzole administration in preventing the delayed impact of acute stress in the amygdala. Hence, in the present study we examined if administration of riluzole in drinking water after stress is effective in reversing the delayed behavioral, structural and physiological effects in the amygdala.

## Results

### Oral administration of riluzole after acute stress prevents its delayed anxiogenic effects

A single episode of 2-hour immobilization stress has been reported to cause a delayed anxiogenic effect (16,29,30,35). We, therefore, wanted to first reproduce this result before examining the efficacy of any post-stress pharmacological intervention (Fig.1A). In the elevated plus-maze task, we observed a significant reduction in the percentage of time spent in the open-arm (Fig.1C: *Control*: 61.3 ± 2.9%; *Stress*: 37.1 ± 5.3%, N=16 rats per group) 10 days after acute immobilization stress. This was also paralleled by a decrease in the percentage of entries into the open-arm (Fig.1D: *Control*: 68.0 ± 1.9%; *Stress*: 41.6 ± 5.0%). Together, these changes were reflected as a significant increase in the anxiety index (*Materials and Methods*) in the stressed rats (Fig.1E: *Control*: 0.35 ± 0.02; *Stress*: 0.61 ± 0.05). Thus, in agreement with previous reports, a single 2-hour exposure to immobilization stress led to a significant enhancement in anxiety-like behavior in the elevated plus-maze 10 days later.

**Figure 1:**
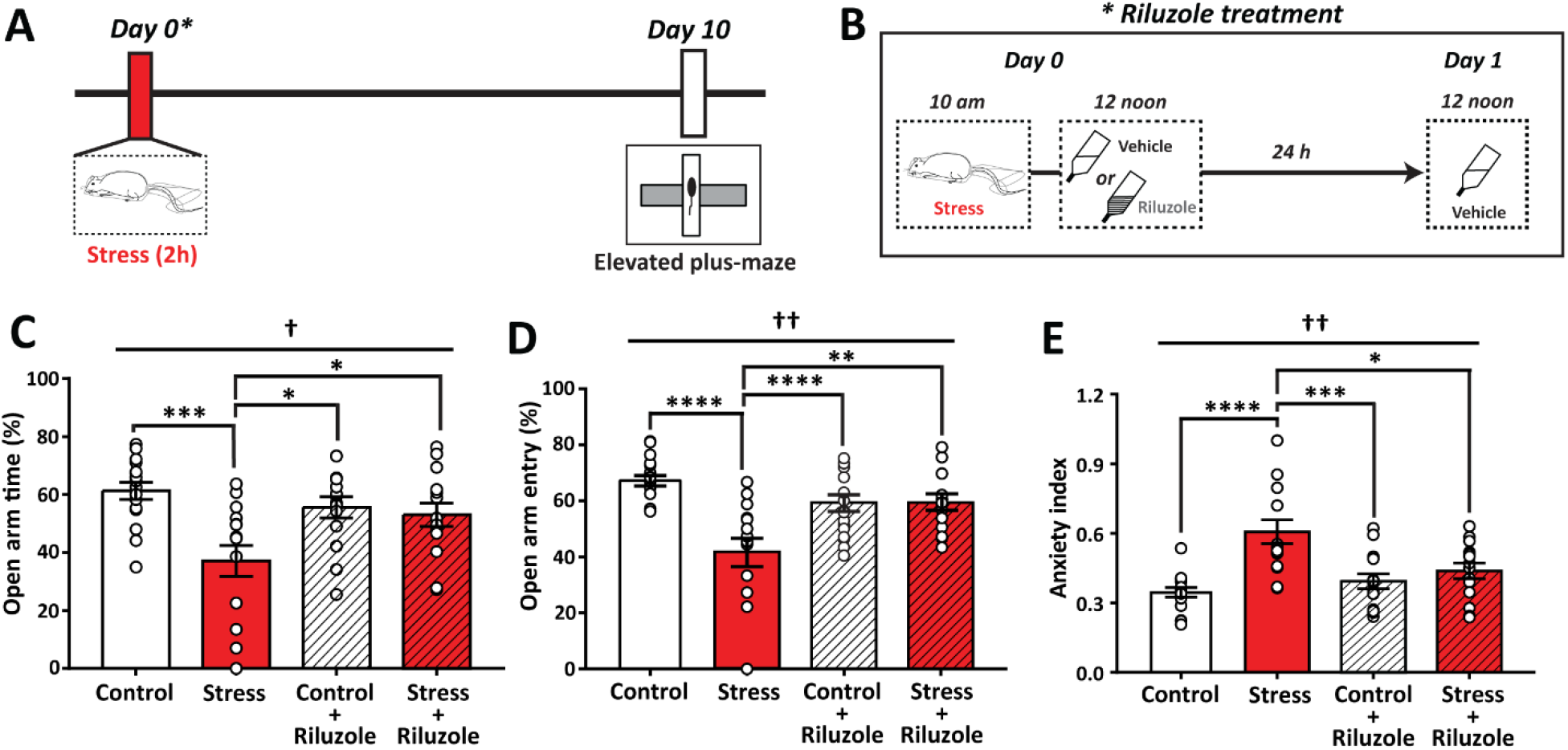
Riluzole in drinking water after stress prevents the delayed increase in anxiety-like behavior. (A) Experimental design depicting the timeline of single episode of immobilization stress (2 hours), followed by riluzole treatment in drinking water for 24 h (*asterisk in A*). After 10 days, anxiety-like behavior was assessed using elevated plus maze. (B) Riluzole is administered on day 0 after the end of 2h stress and continued for 24 h into day 1. On day 1, the drug is replaced by regular drinking water. Summary graph showing % time spent in open arm (C) and %entries into open arm (D) on the elevated plus-maze and overall anxiety index (E) on day 10. Two way ANOVA, post-hoc Tukey’s multiple comparisons test, *p<0.05, **p<0.01, ***p<0.001, ****p<0.0001. †p<0.05, ††p<0.001 in ‘interaction’ between factors stress and riluzole.

Next, a separate cohort of animals were treated with riluzole for 24 hours in the drinking water, at a dosage that was previously shown to be effective in preventing the impact of stress in the rat hippocampus and PFC (31,33,36). Access to regular drinking water was resumed at the end of riluzole treatment (i.e., Day 1, Fig.1B). This post-stress administration of riluzole increased the percentage of time spent in the open-arm specifically in the stressed rats, but not in the control animals (Fig.1C: *Control + Riluzole*: 55.6 ± 3.7%; *Stress + Riluzole*: 53.1 ± 4.0%, N=14 rats per group). Importantly, riluzole administration reversed the stress-induced reduction in open-arm time to levels that were significantly higher than those seen in the stressed animals that did not receive this post-stress treatment. A similar reversal was also seen 10 days later in the open-arm entry in the stressed animals treated with riluzole, without any effects in the control animals (Fig.1D: *Control + Riluzole*: 65.8 ± 2.9%; *Stress + Riluzole*: 59.4 ± 3.1%). As a result, the anxiety index for riluzole-treated stressed rats was comparable to the unstressed control rats that did not receive the drug (Fig.1E: *Control + Riluzole*: 0.39 ± 0.03%; *Stress + Riluzole*: 0.44 ± 0.03%). Significant interaction between the factors “stress” and “riluzole” reflect corrective effects of riluzole on the delayed increase in anxiety-like behavior in stressed rats.

### Oral administration of riluzole after acute stress reverses delayed spinogenesis in the basolateral amygdala

The delayed increase in anxiety-like behavior after acute stress is accompanied by morphological changes in the principal neurons of the rat basolateral amygdala (BLA) (16,30,35). Specifically, the same acute stress exposure causes an increase in the density of spines along the primary apical dendrites of BLA neurons, also 10 days later (16). We, therefore, examined if post-stress riluzole treatment also prevents the delayed BLA spinogenesis 10 days after acute stress (Fig.2A). Hence, we prepared coronal brain sections from all four experimental groups 10 days later and labelled BLA principal neurons with Alexa Fluor-488 for quantification of dendritic spine-density (Fig.2A,B). The total number of spines along the primary apical dendrite, up to a distance of 80 μm from the origin of the dendrite, were counted (*Materials and Methods*). Consistent with previous reports, we found a significant increase in the total number of spines, 10 days after acute stress, compared to controls (Fig.2C, D; Total number of spines in 80 μm, *Control*: 184.1 ± 8.18, N=19 rats; *Stress*: 238.4 ± 12.84, N=15 rats). Riluzole administered, for 24 hours, after the end of 2-hour stress reversed this delayed spinogenesis in the BLA (Fig.2C, D; *Control + Riluzole*: 191.6 ± 10.05, N=14 rats; *Stress + Riluzole*: 172.3 ± 12.89, N=12 rats). Next, we quantified the distribution of the spines in 8 consecutive 10-μm dendritic segments along the length of the primary dendrite (adding up to a total length of 80 μm). Here too, stress led to a significant increase in the number of spines along most of the dendritic segments. Similar patterns, however, were not evident in the riluzole-treated stress group (Fig.2E: *Control*: n=44 dendrites, N=19 rats, *Stress*: n=40 dendrites, N=15 rats, *Control + Riluzole*: n=50 dendrites, N=14 rats, *Stress + Riluzole*: n=46 dendrites, N=12 rats).

**Figure 2:**
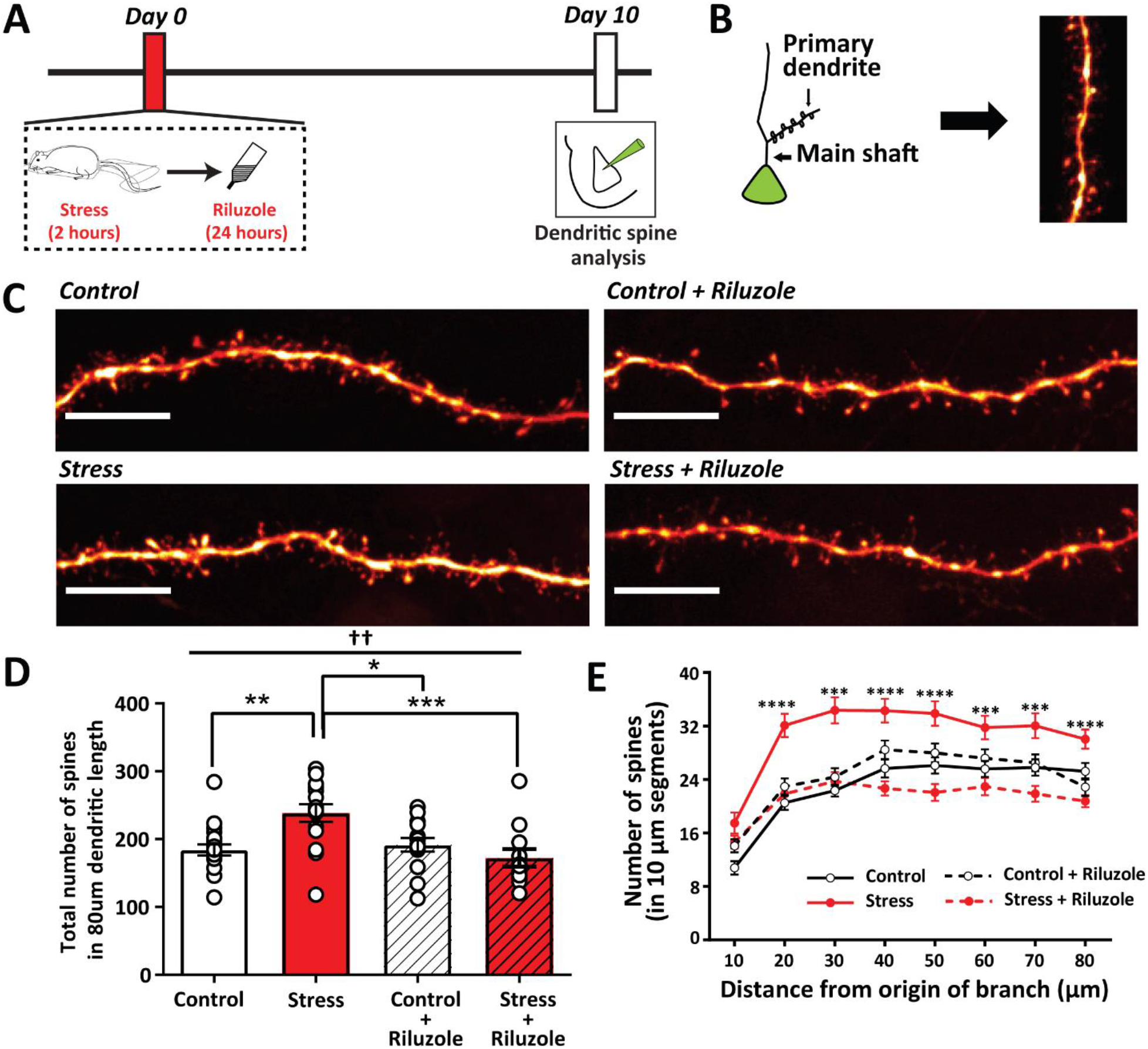
Riluzole in drinking water after stress prevents the delayed increase in spine density on BLA principal neurons. (A) Experimental design. Rats were subjected to a 2 h stress followed by 24 h riluzole treatment on day 0. Iontophoretic dye-labelling carried out on individual BLA principal neurons on day 10. (B) Spines were quantified only on the primary dendrites emanating from the main shaft of the principal neuron. (C) Representative confocal images of dendritic segments and spines from BLA neurons in vehicle and riluzole treated animals (Scale=10μm). (D) Summary bar graph showing total number of spines in 80μm length of primary dendrite. Ordinary Two-way ANOVA, post-hoc Tukey’s multiple comparisons test, *p<0.05, **p<0.01, ***p<0.001, ††p<0.01 in ‘interaction’ between factors stress and riluzole. (E) Plot showing number of spines in successive 10μm segments along the length of a primary dendrite. Repeated measures two-way ANOVA, post-hoc Tukey’s multiple comparisons test for “stress” vs “stress+riluzole”, ***p<0.001, ****p<0.0001.

### Oral administration of riluzole after acute stress prevents delayed increase in synaptic excitability in basolateral amygdala neurons

Dendritic spines form the structural substrate for excitatory synaptic transmission in the rodent brain. The delayed effects of a single episode of acute stress in the BLA are manifested not only as increased spine-density, but also as physiological strengthening through enhanced excitatory synaptic transmission (17,37). Since oral administration of riluzole after acute stress prevents stress-induced spinogenesis, would the same riluzole treatment also block the physiological effects at excitatory synapses?

To address this question, we used whole-cell voltage-clamp recordings in brain slices (*Materials and Methods*) to monitor miniature excitatory postsynaptic currents (mEPSCs) in BLA principal neurons 10 days after a single exposure to 2-hour acute stress (Fig.3A). These BLA neurons typically exhibit accommodating action potential firing in response to depolarizing current injections (Fig.3B). As reported earlier (17,37), stress triggered a delayed increase in the average instantaneous frequency of mEPSCs (Fig.3C, D: *Control*: 1.53 ± 0.17 Hz, n=8 cells, N=6 rats; *Stress*: 2.41 ± 0.25 Hz, n=13 cells, N=6 rats). However, the same stress failed to elicit any increase in mEPSC frequency after oral administration of riluzole and this reversal of stress-induced enhancement of mEPSC frequency was statistically significant (Fig.3C, D: *Control + Riluzole*: 1.25 ± 0.17 Hz, n=9 cells, N=6 rats; *Stress + Riluzole*: 1.49 ± 0.20 Hz, n=12 cells, N=6 rats). The stress-induced increase in mEPSC frequency was also reflected in the cumulative frequency distribution of the interevent intervals (Fig.3E). The distribution curve is shifted to the left in stressed animals, implying a larger number of events with lesser interevent intervals following stress, which is reflected as enhanced frequency. This shift is prevented with riluzole treatment after stress. We did not observe any effects of stress on the average amplitude of these miniature events (Fig.3F: *Control*: 19.36 ± 0.99 pA, *Stress*: 20.18 ± 0.73 pA, *Control + Riluzole*: 19.62 ± 1.18 pA, *Stress + Riluzole*: 19.51 ± 0.69 pA).

**Figure 3:**
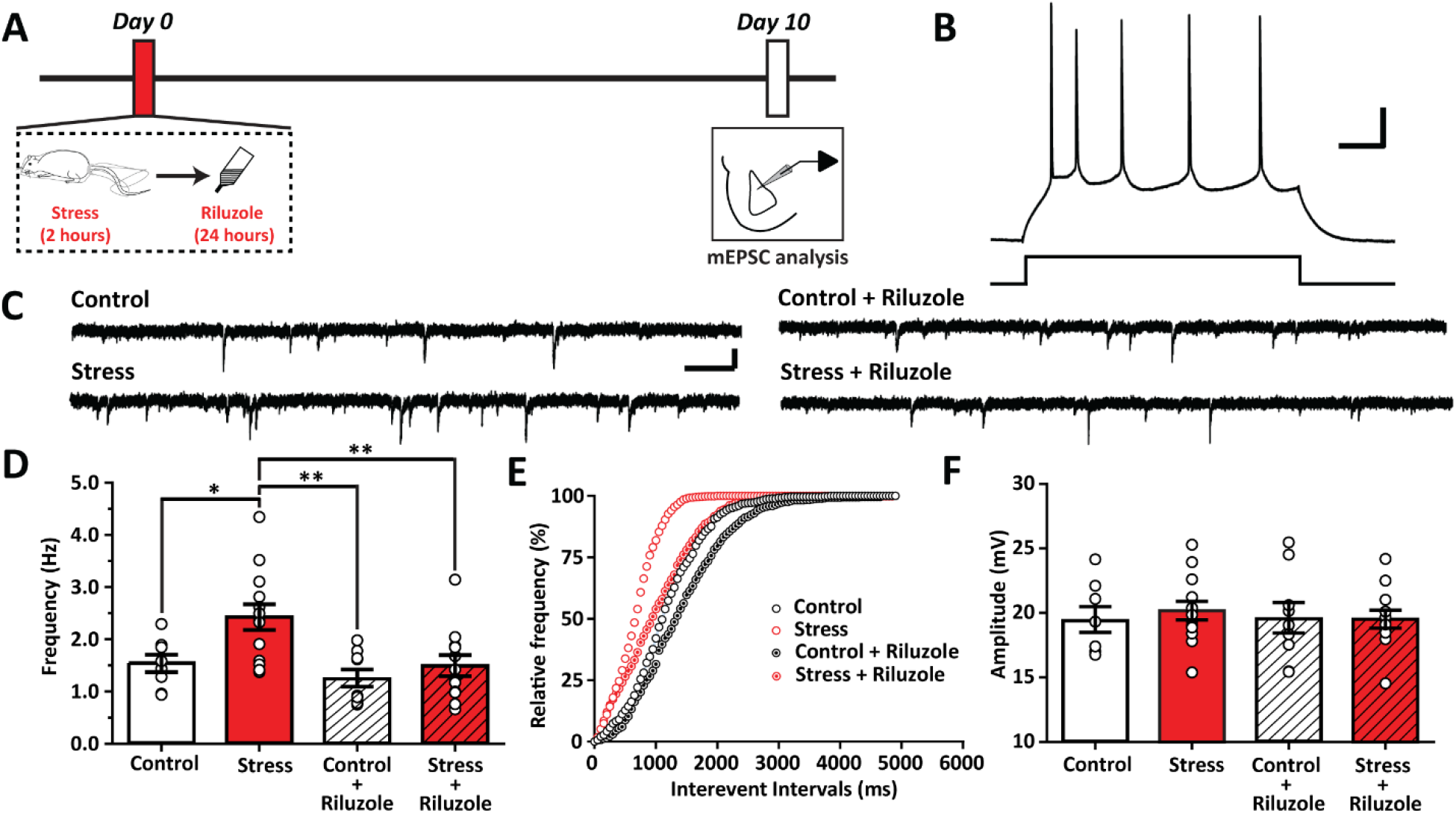
Post-stress riluzole administration in drinking water prevents the delayed increase in frequency of mEPSCs on BLA principal neurons. (A) Experimental design. Rats were subjected to a 2 h stress followed by 24 h riluzole treatment on day 0. Whole cell patch clamp recordings of mEPSCs were carried out on BLA principal neurons on day 10. (B) Representative trace of an accommodating action potential recorded from a BLA principal neuron in response to a depolarizing current injection (Scale: 25mV, 100ms). (C) Representative traces of mEPSCs from BLA principal neurons in vehicle and riluzole treated animals (Scale: 20 pA, 0.5 s). (D) Summary graph showing frequency of mEPSCs on day 10. Two way ANOVA, post-hoc Sidak’s multiple comparisons test, *p<0.05, **p<0.01. (E) Cumulative frequency distribution of mEPSCs in vehicle and riluzole treated groups. (F) Graph showing average amplitude of mEPSCs in all groups.

## Discussion

Here we tested if suppressing the elevation of extracellular glutamate levels, after stress exposure (13,14,37), prevents its delayed impact on amygdalar structure and function in rats. Specifically, we analyzed if systemic riluzole administration in the drinking water, immediately after acute immobilization stress, reversed the strengthening of the morphological and physiological basis of synaptic connectivity in the BLA, as well as enhanced anxiety-like behavior, 10 days later. This is in contrast to a majority of earlier research that was aimed directly at enhancing GABAergic transmission or using corticosterone to reverse stress-induced effects (29,30,35,38-43). To this end, we coupled this strategy with a rat model of acute stress that offered a time window to intervene, after stress exposure, but well before its delayed behavioral, electrophysiological and morphological effects are manifested. In agreement with previous reports, we observed an increase in anxiety-like behavior in stressed rats on the elevated plus-maze, 10 days later. This was prevented by a 24-hour administration of riluzole in drinking water, beginning immediately after the end of the 2-hour immobilization stress. The enhanced anxiety was also accompanied by stress-induced enhancement in dendritic spine-density on BLA pyramidal-like neurons 10 days later. Again, the same post-stress riluzole treatment for 24 hours prevented this delayed amygdalar spinogenesis. Finally, this stress-induced increase in spine-density was paralleled by a physiological strengthening of excitatory synaptic transmission as evidenced by an increase in the frequency of mEPSCs 10 days after acute stress. This too was reversed by the same post-stress riluzole treatment.

The findings reported here add new dimensions to previous studies on the efficacy of riluzole in countering the detrimental effects of stress. First, while earlier work focused on stress-induced changes in the hippocampus and PFC (31,32,44), this is the first such report on how riluzole is effective in reversing the opposite effects of stress in the amygdala across biological scales spanning behavioral and synaptic changes (Fig.4). Second, we explored the delayed impact of acute stress as opposed to models of chronic or repeated stress in earlier studies. For instance, riluzole administration in a rat model of repeated stress helped rescue anhedonia and helplessness behavior induced by chronic unpredictable stress (31). This same treatment also prevented stress-induced alterations in glial metabolism in the PFC (31). Notably, those earlier analyses that used chronic stress paradigms, combined them with chronic riluzole treatments that overlapped with the entire duration of the chronic stress (31-33). In this context, it is interesting to note that the duration of riluzole treatment itself can give rise to strikingly different effects. For example, in a previous study (33), riluzole administration in the drinking water was initiated before stress and continued for the entire duration of 10 days of chronic immobilization stress (2h/day). This prevented the loss of dendritic spines on hippocampal CA1 pyramidal neurons caused by chronic stress, though the drug by itself had no effect. On the other hand, the same chronic riluzole treatment in the same animals failed to prevent stress-induced spinogenesis in the BLA. However, chronic riluzole administration alone also led to spinogenesis in the BLA. Thus, the same chronic administration of riluzole that blocked the morphological effect of stress on spines in the hippocampus, instead mimicked its effect on amygdalar spines (33). In contrast, here we find that a single 24-hour treatment with riluzole by itself did not have any morphological or electrophysiological effects on the amygdala. Third, unlike these earlier studies, riluzole was administered after the end of acute stress. While this was effective in reversing the delayed effects 10 days later, it remains to be seen if riluzole administration is necessary immediately after the stress has occurred or whether it will also be effective when used at later time points after acute stress, and how long this post-stress window of opportunity extends. Finally, the administration of riluzole through drinking water offers an advantage over other methods like oral gavage or intraperitoneal injections, which have been shown to enhance corticosterone and interfere with the effects of stress (29,30,35,39,43). This experimental paradigm of oral administration of riluzole after stress also bears greater similarity to a clinically realistic framework.

**Figure 4:**
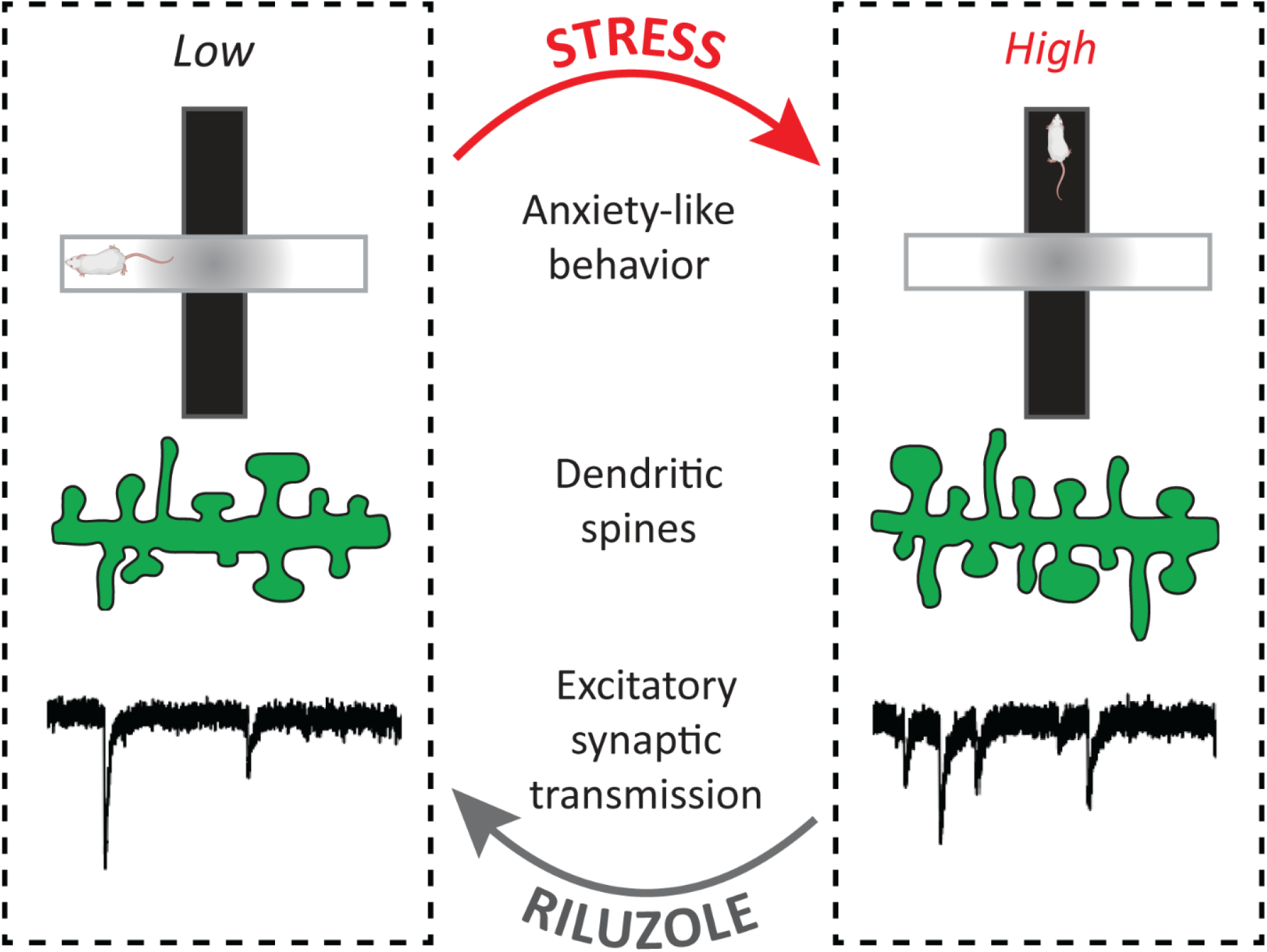
Summary. Acute stress (top) causes a delayed increase in anxiety-like behavior, BLA spine-density and mEPSC frequency. These behavioral and synaptic changes are reversed by post-stress riluzole treatment (bottom).

Riluzole is known to reduce extracellular glutamate levels through its action on several targets. Apart from enhancing glial uptake of glutamate (19,20) and reducing presynaptic glutamate release (22), it has also been reported to act as an antagonist of NMDA and AMPA subtypes of glutamate receptors (18,23). A similar dosage of riluzole was used in previous studies involving the hippocampus and the PFC, (31,33,36), thereby providing a basis for comparing the findings in the BLA reported here. Interestingly, in addition to ALS, riluzole has also been used to treat psychiatric disorders (21,28,45,46). For instance, there is clinical evidence for the antidepressant effects of riluzole (21,31,32,47-50). Further, according to a recent study, riluzole treatment on war veterans resistant to other drugs like SSRIs showed an improvement in hyperarousal, but not in other PTSD symptoms (50). Therefore, the precise mechanisms by which riluzole reverses the delayed effects of stress requires further analyses. It is interesting to note that the astrocytic glutamate transporter GLT-1, which plays a major role in glutamate uptake in the CNS, is potentiated by riluzole (20). The glial glutamate uptake transporters have a higher affinity for riluzole than the other targets (19) enabling their activation at lower concentrations of the drug. Notably, the concentration of riluzole used in our study is comparable to the EC50 of riluzole’s effects on GLT-1. Further, altered GLT-1 expression has been observed in brains of patients with major depressive disorder (MDD) (51-53). Thus, together these findings underscore the need for further investigations into modulation of astrocytic glutamate uptake as a potential therapeutic target for stress disorders.

## Materials and methods

### Animals and stress protocol

All experiments were performed on male Wistar rats (postnatal 55-60 days of age) maintained on a 14h:10h light-dark cycle, with *ad libitum* food and water. All procedures related to animal care and usage were approved by Institutional Ethics Committee at the National Centre for Biological Sciences, India. Rats were subjected to a single episode of immobilization stress on Day 0 (Fig.1), between 10AM to 12PM, without any access to food and water. At the end of the stress episode, the rats were returned to their home cages and remained unperturbed until day 10. All experiments were performed on day 10. The control rats were not subjected to any such stress episodes and were housed in separate rooms.

### Drug administration

Riluzole (Sigma, St. Louis, MO) was administered to the rats for 24 hours in their drinking water (4mg/kg body weight/day), as established previously (31,33,36). Rats subjected to immobilization stress had access to riluzole in their home cage soon after the termination of the immobilization episode on Day 0 (Fig.1). The control rats from the same cohort also received riluzole for 24h on the same day. This was replaced by regular drinking water following the 24h riluzole treatment. Riluzole solution (∼30μg/ml, based on average water consumption/cage/day) was prepared by dissolving the drug in drinking water by constant stirring overnight ∼6-10h in a light protected container.

### Elevated Plus Maze

Rats were subjected to elevated plus maze (EPM) test 10 days after the exposure to single episode of immobilization stress (30). Control rats from the same cohort were subjected to EPM on the same day. Following ∼20 minutes of habituation in a holding room, rats were placed in the center of the EPM with a free choice to explore open arms (∼75 lx) and closed arms (∼0 lx) for 5 minutes. All trials were video recorded and analyzed offline by the experimenter blind to the assigned group of the animals. The time spent in open arms, number of open arm entries and anxiety index was calculated as described (30), using the following equation:

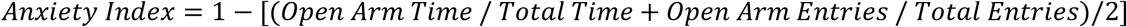

### Intracellular dye fills

10 days after AIS exposure, the stressed and age-matched control animals (∼post-natal day 70) were anesthetized with an overdose of ketamine and xylazine (3:1, 0.9ml ketamine; 0.3ml xylazine) and perfused transcardially with 50 ml of 0.1M phosphate buffer (PB) followed by 100 ml of fixative solution containing 4% 0.1M sodium phosphate-buffered paraformaldehyde (pH 7.4) (at 4°C). Subsequently, the brains were gently removed from the skull and post fixation was done for 3 hours in the above-mentioned fixative (ice cold) on a rocker (at 4°C). Coronal sections of the brain containing the amygdala and dorsal hippocampus were obtained using VT1200S vibratome (Leica Biosystems, Germany) and stored in 0.1M PB for intracellular fills (section thickness: 100 μm).

Neurons were dye labelled using methodology described previously (54). Briefly, the sections were placed in cold PB and the region of interest was identified with an infrared differential interference contrast (DIC)/epifluorescent microscope (SliceScope, Scientifica, UK) using a 4X air objective (Olympus, Japan). The large soma of principal neurons (diameter∼15 micron) were identified within the basolateral amygdala using a 40X water immersion objective (Olympus, Japan). The cell bodies were then impaled with sharp glass micropipette containing 10 mM Alexa Fluor 488 hydrazide, sodium salt solution (Thermo Fisher Scientific, CA, USA). Iontophoretic dye injection into the cells was performed by applying a 0.5s negative current pulse (1 Hz) using Master8 (A.M.P.I. Systems, Israel) until the distal dendrites were sufficiently filled (∼15 min). The sections were then placed in cold 4% paraformaldehyde–PB for 30 minutes followed by DAPI (Sigma, St. Louis, MO) staining for 15 minutes. The sections were then mounted in ProlongGold antifade reagent (Thermo Fisher Scientific, CA, USA).

### Image visualization and analysis

Image acquisition was done using confocal laser scanning microscopy on Olympus Fluoview3000 (Olympus, Japan). Primary dendritic segments of dye-filled neurons were visualized under a 60X oil objective and confocal stacks were acquired at a Z-step size of 0.35 μm (Nyquist sampling criteria) capturing the dendrite origin point and a dendrite length of ∼100 micron. The confocal stacks of individual primary dendrites were used to quantify the number of spines using filaments plug-in on IMARIS (Oxford instruments, Bitplane, Switzerland).

### Electrophysiology

Rats were deeply anaesthetized with halothane, decapitated and the brain removed rapidly. The brain was dissected in ice-cold artificial cerebrospinal fluid (aCSF) containing (in mM): NaCl, 86; glucose, 25; sucrose, 75; NaHCO_3_, 25; NaH_2_PO_4_, 1.2; KCl, 2.5; CaCl_2_, 0.5; and MgCl_2_, 7 (pH 7.4, 320 mOsm) and 400μm coronal sections were cut using VT1200S vibratome (Leica Biosystems, Germany). The sections were allowed to recover for at least 30 minutes in recording aCSF (in mM): NaCl, 124; glucose, 20; NaHCO_3_, 25; NaH_2_PO_4_, 1.2; KCl, 2.5; CaCl_2_, 2; and MgCl_2_, 1 (pH 7.4, 320 mOsm) which was equilibrated with 95% O_2_/5% CO_2_ before being transferred to a submerged chamber for whole-cell recordings from BLA principal neurons under IR-DIC visualization (BX51WI, Olympus).

Miniature excitatory postsynaptic currents (mEPSCs) were recorded from principal neurons in the BLA using whole-cell pipettes (3–5 MΩ) filled with (in mM): CsOH, 110; D-gluconic acid, 110; CsCl, 20; HEPES, 10; NaCl, 4; QX-314, 5; phosphocreatine, 10; Mg-ATP, 4; Na_2_-GTP, 0.3; EGTA 0.2 (pH 7.3-7.4, 280-285 mOsm). Recordings were obtained using a HEKA EPC10 Plus amplifier (Heka Electronik) filtered at 2.9kHz and digitized at 10kHz. Only cells with membrane potentials lesser than -60mV were included. Recordings were discarded if series resistance (Rs) changed by more than 20% from beginning to end or if Rs exceeded 25MΩ. Cells were held at -70mV and mEPSCs were pharmacologically isolated by adding TTX (0.5μM) and picrotoxin (100μM) to the recording aCSF. Continuous current traces of 5-min duration were analyzed using MiniAnalysis (Synaptosoft, NJ).

### Statistical Analysis

All statistical analyses were performed in GraphPad Prism (GraphPad software Inc., La Jolla, CA, USA.). Results are expressed as mean ± SEM. Specific details of tests are mentioned in figure legends.

## Acknowledgments

We thank the Central Imaging and Flow Cytometry Facility, Bangalore Life Science Cluster. We also thank Biorender.com for help in providing illustration. We thank Dr. Siddhartha Datta for his immense contribution towards this project before his untimely demise in 2021. This work was supported by funds from the Tata Institute of Fundamental Research, Department of Atomic Energy and Department of Biotechnology, Government of India (S.D., Z.R., S.N., S.C.). S.D. was also supported by the SERB-National Post Doctoral Fellowship by the Department of Science & Technology, Government of India.

## Data availability statement

All data are included in the manuscript.

